# Using Oyster Genomics to Understand Human Health

**DOI:** 10.64898/2026.11.08.765187

**Authors:** Walker Bargmann, Zhiyuan Chen, Shiying Wang, Matthew Liam Cawood, Akshada Parulekar, Christopher Martucci, Josephine Hoh

## Abstract

As sessile filter feeders that experience continuous environmental exposure, oysters—particularly *Crassostrea virginica*—offer a unique opportunity to understand how environmental exposure alters the genome and influence health. Furthermore, oysters must be sorted by size for market value for premium size and taste. However, it requires intensive labor, specialized equipment (tumblers), and is susceptible to environmental risks like pollution. Therefore, it is desirable to produce uniform oysters without machines. However, limitations in the previous reference genome [2] have constrained our understanding of how genes are organized and maintained in oyster populations. To this end, we harvested 240 adult oysters in same location, 120 extremely large size and 120 extremely small, controlled with all confounding factors such as age and water temperature. We perform a genome wide associate study (GWAS) using next generation whole genome sequencing, aiming to discover genomic regions associated with oyster size. Using PacBio HiFi, Illumina, and Hi-C technologies, we generated a 99.7% complete chromosome-level genome assembly for *C. virginica* (NCBI RefSeq GCF_053477285.1) [75].

We identified a ∼13 Mb region on chromosome 1 uniquely expanded in *C. virginica* and enriched for innate immune and stress-response genes. To evaluate whether these structural features are reflected at the population level, we then conducted a k-mer analysis. This revealed differential sequence patterns by size, including enrichment of bacterial DNA in smaller individuals and host variation at the *RNF213* locus within this region. By highlighting genome structure as a mediator of immune resilience, this work supports the use of oysters as sentinels to advance out understanding of how the dynamic reassembly of the genome due to environmental exposures influence health in human populations.

## Introduction

The adult eastern oyster (*Crassostrea virginica*, hereafter *C. virginica*) is a sessile marine invertebrate native to the Atlantic coast of North America. As a sessile filter feeder, *C. virginica* is subject to persistent variation in salinity [3,4], temperature [5,6], oxygen availability [7,8], and pathogen load [9–11], conditions that are thought to promote lineage-specific expansion of immune and stress-response gene families [12,13]. The genomic mechanisms controlling the dynamic homeostatic processes have yet to be fully revealed in the current literature. Understanding such the physiological responses to environmental stressors has direct implications for better understanding how modern emerging environmental threats can impact human health [9–11, 14]

One such area of interest is innate immunity, the first line of defense against pathogens and foreign threats across taxa [15]. Unlike adaptive immune responses, which are characterized by specificity and memory, innate immune systems rely on broad recognition strategies and rapid activation. Oysters rely solely on innate immunity [16, 17], while humans possess both innate and adaptive immunity mechanisms [18]. Innate immunity plays a critical role in detecting environmental hazards, initiating inflammatory responses, and shaping susceptibility to downstream disease [12, 13, 19]. However, studying how sustained environmental exposures influence innate immune capacity in human populations is inherently challenging due to ethical constraints, heterogeneous exposures, and long latency periods between exposure and health outcomes.

Thus, oysters offer a promising role to serve as a sentinel species to better understand the biological mechanisms by which chronic environmental stress shapes immune capacity. A sentinel species is defined as a non-human organism whose biological responses provide early or amplified indicators of environmental hazards that may pose risks to population health [20].

Such organisms often experience more intense or continuous exposure than humans and lack the behavioral or technological means to mitigate these exposures, allowing biological responses to environmental stress to be observed with greater clarity [20].

However, previous genome assemblies like *C_virginica-3.0* [2] limits such understanding and analysis due to an incomplete reference genome. With haplotig redundancy and chromosomal misassemblies [21], any claims based on gene diversification and cross-species comparisons are invalid [22]. Furthermore, any extrapolation to contexts in human population health cannot be made. As such, this study resolves the issues of the oyster genome assembly to allow for a high-quality genomic comparison across species using a full reference genome [75]. To identify host genomic variants associated with physiological performance in natural populations, we conducted a genome-wide association study (GWAS) of oyster body size. In *C. virginica*, body size is a commercially and ecologically important trait that reflects cumulative growth, environmental exposure, and overall physiological condition [3–6, 9–11]. Because growth in oysters is shaped by both genetic and environmental factors, size can serve as an integrative phenotype capturing the effects of sustained environmental stress. We collected 240 adult *C. virginica* specimens from coastal Connecticut, selecting individuals representing extreme size phenotypes (Large vs. Small). Using whole-genome sequencing in conjunction with our newly assembled reference genome, we tested for allele frequency differences between groups to identify candidate loci associated with variation in growth and physiological performance.

Comparative analyses included four closely related *Magallana* species (*M. gigas, M. hongkongensis, M. ariakensis,* and *M. angulata*) to evaluate lineage-specific immune gene expansion within the genus. The European flat oyster (*Ostrea edulis*) was included as a more distantly related species to distinguish shared oyster immune features from those unique to *C. virginica*. While comparative genomics can identify lineage-specific structural features, it remains unclear whether such regions are actively variable within populations exposed to environmental stress. To address this gap, we complement reference-based analyses with a reference-independent k-mer framework, enabling detection of sequence-level variation across wild oyster populations.

Using this combined approach, we identify extensive intrachromosomal remodeling as a primary driver of genome divergence and characterize a large immune- and stress-enriched genomic region as a prominent feature of *C. virginica*. K-mer analysis shows host variation at the *RNF213* locus, highlighting the adaptability to the genome in contemporary wild populations. These findings position oysters as sentinel systems for elucidating general principles by which genome structure and innate immune capacity respond to sustained environmental exposure. This framework provides a foundation for understanding how environmental exposures may interact with the genome to shape health at the population level.

## Methods

### Developing a Reference Genome

#### Sample Collection and Storage

Adult *C. virginica* oysters were collected from three coastal locations in Connecticut: Sherwood Island Mill Pond (n = 140), Stamford deep-water bed (n = 60), and Westport near-shore bed (n = 40). At each location, specimens were selected to represent extreme size phenotypes (Large and Small groups) while controlling for age by sampling from the same cohort. This stratified sampling design ensured that observed genomic differences between size groups would not be confounded by age or location effects. Mass (g) was recorded for all specimens as the primary size metric, as it demonstrated the greatest reliability across collections. Differences in mean mass between Large and Small groups were assessed using one-way ANOVA to confirm significant phenotypic separation. See table 1 in the results to see mean oyster mass by collection area.

**Table 1.** Summarized dimensions of small and large oysters.

| **Oyster Group** | **Specimen Count** | **\*Median Length (mm)** | **\*Median Width (mm)** | **\*Median Height (mm)** | **\*\*Median Mass (g)** |
|---|---|---|---|---|---|
| Large | 88 | 79 | 55 | 24 | 44.0 |
| Small | 86 | 56 | 43 | 16 | 20.4 |

Table summarizes the medians of length, width, and height in millimeters, and mass in grams.
\*Measurements were estimated to the nearest whole number  \*\*Weight taken with CS Series Portable Balance CS200

Live oysters were harvested and immediately placed on ice from transport to the laboratory, where they were subsequently shucked and tissue samples were prepped and flash-frozen via liquid nitrogen. Three samples of mantle tissues and one sample of gut tissue were isolated from each specimen and kept for long-term storage in -80 degree Celsius freezers to preserve the molecular integrity of the DNA.

### DNA Extraction and Quality Control

Genetic material was extracted primarily from mantle tissue samples using the E.Z.N.A. Mollusk and Insect DNA Kit (Omega Bio-tek) with modifications to the manufacturer’s protocol to optimize yield from oyster tissue. To minimize technical variability, extractions were performed in batches of no more than four samples at a time. Sample quality was assessed using two complementary approaches. First, NanoDrop spectrophotometry (Thermo Fisher Scientific) was used to evaluate total nucleic acid concentration and purity, with acceptable thresholds defined as a 260/280 ratio of 1.8–2.1 and a 260/230 ratio > 1.8. Second, double-stranded DNA (dsDNA) concentration was quantified using a PicoGreen-based fluorometric assay (SpectraMax/PicoDrop), which selectively measures dsDNA and avoids overestimation from RNA or other contaminants inherent to spectrophotometric methods. Samples with dsDNA concentrations ≥ 10.79 ng/µL and acceptable purity ratios were retained for downstream sequencing and genotyping analyses

### Genetic Sequencing, Genome Assembly, and Annotation

Genome assembly was performed by integrating multiple complementary sequencing technologies to generate a chromosome-level reference genome. High-quality DNA was fragmented and converted into sequencing libraries compatible with each platform, ensuring representative sampling across the genome. Whole-genome shotgun sequencing was performed on the Illumina NovaSeq X platform, producing millions of short reads that provided broad, non-targeted coverage and enabled detection of single nucleotide polymorphisms (SNPs) and other small variants. PacBio HiFi long-read sequencing was conducted on the Revio system using Single Molecule Real-Time (SMRT) sequencing, generating long reads thousands of bases in length. These reads were critical for resolving complex genomic regions, including repetitive elements and structural variants that are difficult to assemble using short-read data alone. Hi-C chromatin conformation capture was performed using the Arima-HiC Kit to capture three-dimensional genome organization.

Hi-C sequencing identifies DNA fragments that are physically proximal within the nucleus, enabling the generation of chromosomal contact maps used to order, orient, and scaffold contigs into chromosome-scale assemblies. Hi-C libraries were sequenced on the Illumina NovaSeq X platform. The integration of short-read sequencing, long-read sequencing, and Hi-C contact maps enabled the generation of a highly contiguous, chromosome-level genome assembly with improved completeness and reduced redundancy relative to the previous reference genome.

Genome annotation was performed to characterize gene content and structural features of the genome. Repeat elements were identified using standard repeat annotation pipelines, revealing that a substantial portion of the genome consists of transposable elements and other repetitive sequences. Gene prediction was conducted using a combination of ab initio and evidence-based approaches, incorporating transcriptomic and protein homology data, resulting in the annotation of 33,259 protein-coding genes. Functional annotation was assigned through sequence homology searches against established protein databases. Additional analyses identified key genomic features, including telomeric repeats and putative centromeric regions. Together, these approaches enabled characterization of genome organization and the spatial relationship between gene-rich and repeat-dense regions, providing a framework for downstream comparative and functional analyses.

### Whole-Genome Sequencing and Variant Identification

Whole-genome sequencing was performed on the Illumina NovaSeq X platform to generate genome-wide sequence data for variant discovery. Sequencing data were processed to identify genetic variation across samples relative to the newly assembled *C. virginica* reference genome (NCBI RefSeq GCF_053477285.1) [75]. Candidate single nucleotide polymorphisms (SNPs) were identified from these data for downstream analysis.

Several candidate SNPs were selected for further evaluation based on preliminary computational analyses, including variants located within conserved haplotype blocks identified in the reference genome. One such SNP, located on HiC_scaffold_1 at position 35,698,255 (A>G), was prioritized for downstream genotyping and association testing. This variant resides within an intronic region of a predicted protein-coding gene (g2168) and was selected based on its statistical association with the size phenotype in preliminary analyses.

### Association Testing

Association between genotype and oyster size phenotype (Large vs. Small) was evaluated using Fisher’s exact test for individual biallelic SNPs. This approach was selected due to the modest sample size and categorical classification of the phenotype.

For each candidate SNP, 2×2 contingency tables were constructed comparing the presence or absence of the alternate allele between Large and Small oysters. Two-tailed p-values were calculated to assess statistical significance of the association.

For the lead SNP (HiC_scaffold_1:35698255, A>G), genotype counts were tabulated by phenotype group to evaluate the direction and strength of association. This variant demonstrated a statistically significant enrichment of the alternate allele in the Large phenotype group, supporting its role as a candidate locus associated with variation in oyster size.

### Comparative and Functional Analyses of the Genome

Comparative genomic analyses across six other oyster species—*Ostrea edulis, Magallana gigas, M. angulata, M. ariakensis,* and *M. hongkongensis*—using publicly available reference genome assemblies obtained from NCBI genomic databases [2] and prior studies [22]. Whole-genome alignments and synteny analyses were performed to evaluate chromosomal conservation and structural divergence across species. Regions of reduced ortholog conservation were identified as candidate loci of lineage-specific variation. Functional enrichment analyses were conducted to characterize gene content within these regions, with a particular focus on immune and stress-related pathways. To assess transcriptional activity, previously published RNA-seq data from immune-stimulated and control samples were analyses to evaluated gene expression patterns within structurally dynamic genomic regions [23–25].

### Reference-Independent K-mer Analysis

To complement reference-based genomic analyses, we performed a reference-independent k-mer-based association analysis to identify sequence-level variation between groups without reliance on genome alignment. This approach enables detection of variants that are often missed by standard alignment-based methods, including structural rearrangements, repetitive elements, and non-host (e.g., microbial) sequences.

### Data Processing and k-mer Counting

Whole-genome sequencing (WGS) data from 174 samples (88 in the “Large” group and 86 in the “Small” group) were analyzed. Sequencing reads were processed using *kmtricks* [70] to efficiently count k-mers across all samples. Initial analyses used a k-mer length of 31, with additional analyses conducted at *k* = 25 and *k* = 21 to assess robustness across sequence scales.

### Quality Control

A multi-layered quality control (QC) pipeline was implemented to minimize sequencing artifacts and reduce noise. First, raw sequencing reads underwent quality assessment using FastQC [72], followed by adapter trimming and filtering to retain reads with a minimum length of 50 base pairs. Second, k-mers within each sample were filtered to retain only those with a minimum count of ≥2, removing low-frequency k-mers likely arising from sequencing error. Third, k-mers were filtered across samples to retain only those present in at least five samples, reducing the influence of rare or sample-specific artifacts and lowering the multiple testing burden.

### Association Testing

Differential k-mer frequency between groups was assessed using *kmdiff* [71], which applies a likelihood ratio test within a Poisson regression framework. Analyses were conducted both with and without adjustment for potential confounders, including sequencing batch, sequencing depth, and principal components (PCs) derived from k-mer data to account for population structure.

Multiple testing correction was performed using both Bonferroni correction (threshold of 8.5e-13) and the Benjamini–Hochberg false discovery rate (FDR) procedure [73]. Results across different k-mer lengths were compared to evaluate consistency of detected signals.

### Validation and Sensitivity Analyses

To assess robustness, sensitivity analyses included adjustment for covariates, removal of outlier samples identified through principal component analysis [76], and transformation of k-mer counts to binary presence/absence variables. Independent validation was conducted using a subset of 17 samples (8 Large, 9 Small), with statistical significance evaluated using Fisher’s exact test.

### Local Assembly and Sequence Annotation

To characterize significant k-mers, reads containing these sequences were extracted and subjected to de novo local assembly using long-read PacBio data. Resulting contigs were aligned to the *C. virginica* reference genome using minimap2 [74]. Sequences that did not map to the reference genome were further analyzed using BLAST against multiple databases, including the oyster reference genome and the NCBI nucleotide and protein databases [75], to infer potential sequence origin.

## Results

### Developing a Reference Genome

#### Developing a New High-Quality Reference Genome

We found the final high-quality genome assembly of *C. virginica* contained 10 chromosomes with a length of 529.0 Mb, with 99.7% completeness. This assembly is more complete with fewer redundancies (BUSCO duplication reduced from 18.7% to 1.1%) than the previous reference genome, *C_virginica-3.0* [2]. Additionally, our genome corrects misassemblies in the previous reference genome, namely the fragmentation of chromosome 1, fusion of chromosome 5 and 7, and the misjoining of chromosomes 3, 5, and 10. Quality assessments confirmed high completeness and integrity, with near-complete gene representation and strong chromosome-scale organization supported by Hi-C mapping.

**Table 2.** Mean masses (g) of small and large oysters by collection location.

| **Location (number of specimens)** | **Mean Mass (g) – Large Oysters** | **Mean Mass (g) – Small Oysters** | **Difference in Means (g): (Large – Small)** |
|---|---|---|---|
| Mill Pond (140) | 45.56 | 18.51 | 27.37 |
| Stamford (60) | 62.95 | 31.80 | 31.15 |
| Westport (40) | 47.83 | 28.19 | 19.64 |

Column one shows the location of collection and number of specimens collected; column two shows the mean mass by location of large oysters; column 3 shows mean mass of small oysters by collection location; column four shows the difference in means between small and large oysters by collection location.

#### Phenotype Validation

Mass differed significantly between Large and Small phenotypic groups across all three collection locations (Table 1). At Mill Pond (n = 140), the largest collection site, mean mass was 45.56 g for Large oysters and 18.51 g for Small oysters, representing a 2.5-fold difference between groups. One-way ANOVA confirmed significant between-group variance in mass (F = 858.79, p = 4.08 × 10⁻⁶¹), validating the use of these extreme phenotypes for downstream genetic association analyses.

#### Identification of a Size-Associated SNP on Chromosome 1

Analysis of 18 specimens (8 Large, 10 Small) from the Stamford collection identified a single nucleotide polymorphism (SNP) at position 35,698,255 on HiC_scaffold_1 (corresponding to Chromosome 1 in the assembled reference genome) with a significant allele frequency difference between Large and Small oysters (Table 3). The reference allele (A) and alternate allele (G) showed markedly different distributions between phenotypic groups. The alternate (G) allele was present in 7 of 8 Large oysters (87.5%), including 6 homozygous (G/G) and 1 heterozygous (A/G) individual. In contrast, the G allele was completely absent from all 10 Small oysters tested, all of which were homozygous for the reference allele (A/A). Fisher’s exact test confirmed a statistically significant association between the G allele and the Large phenotype (p = 0.0002514).

This SNP resides within an intronic region of gene *g2168*, a predicted protein-coding gene with no currently annotated function. Variant quality metrics were favorable, including a mapping quality score (MQ) of 47.21, quality-by-depth (QD) of 29.57, and low strand bias (FS = 3.572), indicating a high-confidence variant.

**Table 3.** Summary of Comparative Analysis of SNP 35698255.

| **Group** | **No G Allele** | **Has G Allele** | **Total** |
|---|---|---|---|
| Large | 1 | 7 | 8 |
| Small | 10 | 0 | 10 |

This contingency table for Fisher’s Exact Test yielded p=0.0002514, indicating this SNP is a strong candidate marker for size in *C. virginica*.

#### Localization Within the Immune-Enriched Genomic Neighborhood

Notably, HiC_scaffold_1:35,698,255 falls within the ∼13 Mb region on Chromosome 1 identified in our comparative genomic analyses as enriched for innate immune and stress-response genes (see Results: Developing a Reference Genome). This region contains 908 genes and is characterized by lineage-specific expansion of immune-related gene families in *C. virginica* relative to *M. gigas*. While the identified SNP itself is non-coding, its location within this structurally dynamic genomic neighborhood—which also includes RNF213 and other immune-associated loci—suggests potential linkage with functional variation in nearby genes involved in growth, immune function, or stress physiology.

#### Genome Annotation Reveals Structured Organization of Genes and Repetitive Elements

The annotation of *C. virginica* genome characterized its sequence composition and gene organization. The genome contained 33,259 protein-coding regions, which have provided a comprehensive framework for downstream analyses. A substantial portion of the genome contains repeat elements that are unevenly distributed across chromosomes. Notably, we found gene density to be inversely correlated with repeat content, and gene-rich regions to be distinct from these repeat-dense regions. This spatial organization is consistent with other eukaryotic genomes and suggests a structured genomic landscape in which functional genes are partitioned from highly repetitive sequences [26].

#### Comparative Genomic Analysis of Chromosomal Architecture Across Species

To assess the *C. virginica* genome assembly in evolutionary context, we conducted comparative chromosome-level analyses against five species, including *O. edulis* (xbOstEdul1.1), *M. gigas* (xbMagGiga1.1), *M. angulata* (ASM2561291v2), *M. ariakensis* (ASM4011448v1)and *M. hongkongensis* (ASM1577677v1). We found that all ten chromosomes corresponded to homologous chromosomes in a one-to-one fashion across the five species. While overall global structures seem to be conserved across species, divergence is revealed at finer genomic scales.

Rather than large-scale chromosomal reshuffling, most structural differences are driven by intrachromosomal remodeling, such as local rearrangements, insertions, and inversions that are unevenly distributed. Additionally, comparative analyses against the *Magallana* clade showe that regions of reduced ortholog conservation regions were enriched in immune and stress-related genes within *C. virginica.* This pattern suggests that lineage-specific adaptations are concentrated within discrete genomic regions and lend perspective to our understanding of how gene families can reorganize in response to environmental pressures.

#### Immune Functional Enrichment and Localized Genomic Expansions Highlight Dynamic Architecture

Further comparative analyses were conducted to investigate the composition of dynamic genomic regions. We selected *M. gigas* for comparison as the species is distinct from *C. virginica* while remaining in close phylogenetic relatedness, allowing for an evaluation of genomic regions with reduced ortholog conservation across species. These regions were enriched for genes involved in innate immune responses and environmental stress pathways, including pathogen defense and cellular stress signaling. Further examination of the non-conserved regions showed evidence of localized gene family expansion with immune and stress-related genes occurring in dense clusters. As these expansions are not uniformly conserved across other oyster species, this suggests a genomic adaptation to species-specific environmental pressures rather than shared ancestral features. In other words, our evaluation of regional genomic differences between *C. virginica* and *M. gigas* suggest that the non-conserved orthologs and expansions are not a result of functional randomness, but rather the dynamic nature of the genome and its relationship to gene-environment interactions.

This is pattern most prominently revealed in a 13 Mb region on chromosome 1 containing 908 genes representing the largest contiguous divergent region between *C. virginica* and *M. gigas*. Within this region, *C. virginica* had 36 families of genes that had more than four copies, showing structural differences from *M. gigas*. These expansions contribute to the reduced ortholog conservation relative to *M. gigas*, lending further evidence in support of lineage-specific gene duplication underlying structural divergence in this region. 268 genes within the 13 Mb region were most notably associated with immune and stress-related response pathways, particularly cytokine-mediated signaling, a key component of immune communication. 87 genes showed at least a 2-fold increase in expression during immune stimulation under 2’3’-cUA treatment. This coordinated regulatory activation in response to immune stimulation suggests that this structurally dynamic genomic region harbors functionally integrated gene networks that contribute to adaptative immune responses under sustained environmental stress.

#### Reference-Independent k-mer Analysis Identifies Differential Sequence Patterns

To investigate sequence-level variation independent of reference genome alignment, we performed a k-mer-based association analysis across 174 *C. virginica* samples (88 Large, 86 Small). This analysis identified multiple classes of sequence variation differentiating the cohorts, including signals derived from non-reference sequences, host genomic loci, and repetitive elements.

#### Identification of Differentially Abundant k-mers

Initial analyses using *k* = 31 identified two highly significant k-mers under Bonferroni correction (threshold = 8.5 × 10⁻¹³), both enriched in the Small group (p = 1.24 × 10⁻¹³ and 2.08 × 10⁻¹³). These k-mers differed by a single nucleotide and likely represent the same underlying locus. Following implementation of the full quality control pipeline and evaluation across multiple k-mer lengths (*k* = 21, 25, 31), a greater number of significant k-mers were identified under Benjamini-Hochberg false discovery rate (FDR) correction. The summary can be found in the table below.

**Table 4.** k-mer association results across k-mer lengths and correction methods.

| k-mer length | Total k-mers after QC | Correction (threshold) | Number of significant k-mers | Enriched in small oysters | Enriched in large oysters |
|---|---|---|---|---|---|
| 31 | 4.97 × 10<sup>9</sup> | Bonferroni (1.01 × 10<sup>−11</sup>) | 0 | 0 | 0 |
|  | 4.97 × 10<sup>9</sup> | Benjamin-Hochberg (FDR = 0.05) | 0 | 0 | 0 |
| 25 | 4.24 × 10<sup>9</sup> | Bonferroni (1.17 × 10<sup>−11</sup>) | 6 | 6 | 0 |
|  | 4.24 × 10<sup>9</sup> | Benjamin-Hochberg (FDR = 0.05) | 174 | 168 | 6 |
| 21 | 3.65 × 10<sup>9</sup> | Bonferroni (1.36 × 10<sup>−11</sup>) | 0 | 0 | 0 |
|  | 3.65 × 10<sup>9</sup> | Benjamin-Hochberg (FDR = 0.05) | 743 | 686 | 57 |

Results are shown for k = 21, 25, and 31 after quality control filtering. Bonferroni correction applied thresholds of 1.36 × 10<sup>−11</sup> (k=21), 1.17 × 10<sup>−11</sup> (k=25), and 1.01 × 10<sup>−11</sup> (k=31). Benjamini-Hochberg (BH) correction controlled the false discovery rate at FDR = 0.05. Direction of enrichment indicates whether significant k-mers were more abundant in the Small or Large group.

At *k* = 25, 174 k-mers reached significance, with 168 (96.6%) enriched in the Small group and 6 (3.4%) enriched in the Large group. At *k* = 21, 743 k-mers reached significance (686 enriched in Small, 57 in Large). The magnitude of enrichment in the Small group was substantial, with top k-mers exhibiting nearly and 18-fold higher frequency relative to the Large group. The top five k-mers are shown in the table below.

**Table 5.** Top five most significant k-mers identified at k = 25 under Benjamini–Hochberg correction.

| **Rank** | **p-value** | **Small Count** | **Large Count** | **Enrichment (Small/Large)** |
|---|---|---|---|---|
| 1 | 6.21 × 10<sup>−12</sup> | 107 | 6 | 17.83x |
| 2 | 8.90 × 10<sup>−12</sup> | 110 | 7 | 15.71x |
| 3 | 8.90 × 10<sup>−12</sup> | 110 | 7 | 15.71x |
| 4 | 8.90 × 10<sup>−12</sup> | 110 | 7 | 15.71x |
| 5 | 1.05 × 10<sup>−11</sup> | 105 | 8 | 12.13x |

Counts indicate the number of samples in which each k-mer was observed in the Small and Large oyster groups. Enrichment values represent fold-differences in frequency between groups.

#### Robustness to Covariate Adjustment and Sensitivity Analyses

The identified k-mer associations were robust to multiple sensitivity analyses. Inclusion of sequencing batch, sequencing depth, and principal components (PC1–PC10) as covariates did not materially alter statistical significance (adjusted p-values ranging from 1.33 × 10⁻¹¹ to 1.55 × 10⁻¹²). Removal of outlier samples identified through principal component analysis and transformation of k-mer counts to binary presence/absence variables yielded consistent results, indicating that the observed associations were not attributable to technical confounding or population stratification [76].

**Table 6.**
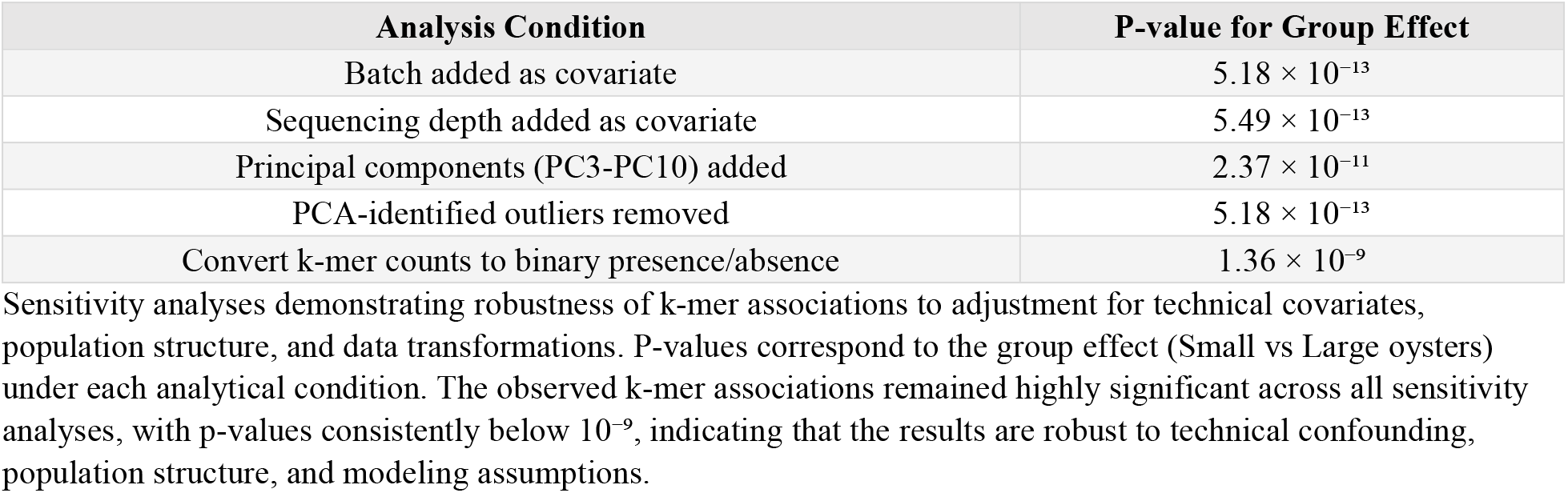
Sensitivity analyses demonstrating robustness of k-mer associations to covariate adjustment and data transformation.

| Analysis Condition | P-value for Group Effect |
|---|---|
| Batch added as covariate | 5.18 × 10<sup>−13</sup> |
| Sequencing depth added as covariate | 5.49 × 10<sup>−13</sup> |
| Principal components (PC3-PC10) added | 2.37 × 10<sup>−11</sup> |
| PCA-identified outliers removed | 5.18 × 10<sup>−13</sup> |
| Convert k-mer counts to binary presence/absence | 1.36 × 10<sup>−9</sup> |

#### Characterization of Non-Reference Sequence Signals

Local assembly of PacBio long reads containing the most significant k-mers produced a 525 bp contig supported by 16 long reads, with read lengths ranging from approximately 4 kb to 29 kb. Alignment of this contig to the *C. virginica* reference genome [75] revealed minimal mapping, with only short, low-coverage alignments observed across fragmented genomic regions. Similar weak alignments were observed across other available oyster reference genomes [2].

BLAST analysis [77] against nucleotide databases did not identify strong matches within curated oyster reference assemblies. However, alignment against whole-genome shotgun (WGS) datasets revealed near full-length matches to sequences from related oyster species [2]. Protein-level BLAST (BLASTx) identified similarity to a hypothetical bacterial protein from the *Spirochaetota* phylum (E-value = 1 × 10⁻²³; 98% query coverage), suggesting that this signal may represent exogenous microbial DNA rather than a canonical oyster genomic sequence [78].

Additional bacterial sequence matches, including those from the *Actinomycetota* phylum [79], were also observed among significant k-mers.

#### Host Genomic Signal Localizes to Chromosome 1

Among the k-mers enriched in the Small group, a subset mapped unambiguously to the *C. virginica* reference genome [75]. These k-mers localized to Chromosome 1, specifically within the *RNF213* gene (E3 ubiquitin-protein ligase mf213-alpha-like; 99.7% nucleotide identity). This locus resides within a region corresponding to the immune-enriched genomic neighborhood identified in the reference genome analysis.

Inspection of PacBio long reads spanning this region indicate that some reads contained both oyster genomic sequence and non-reference sequence segments, consistent with the presence of chimeric reads, potentially arising during library preparation or sequencing.

#### Low-Complexity K-mers Enriched in the Large Phenotype

The k-mers enriched in the Large group were characterized by low sequence complexity and high AT content (68–80%). These k-mers mapped to repetitive genomic regions distributed across multiple chromosomes, including loci overlapping ribonuclease-associated gene clusters.

This pattern is consistent with differential representation of repetitive or low-complexity sequences between cohorts.

#### Limited Validation in Independent Samples

Validation in an independent subset of 17 samples (8 Large, 9 Small) showed directional consistency with the discovery analysis (k-mer present in 3/9 Small vs. 1/8 Large), although this difference did not reach statistical significance (Fisher’s exact test, p = 0.577).

## Discussion

### Genomic Plasticity of Oysters Reflects the Dynamic Gene-Environment Interaction

This study presents a high-quality genome assembly of *C. virginica* that improves upon the previous reference (C_virginica-3.0) by resolving redundancies, correcting misassemblies, and increasing completeness to BUSCO 99.7%. Through gene annotation and comparative analyses with other closely related oyster species, we identified a ∼13 Mb region on chromosome 1 characterized by extensive gene expansion, particularly among genes involved in immune and stress-response pathways. Notably, these expansions were not conserved across *Ostreidae*, suggesting that intrachromosomal rearrangements and localized gene family expansions are driven by species-specific environmental exposures rather than shared evolutionary history. This pattern supports a model in which genome structure itself is shaped by adaptation to environmental stressors.

To further evaluate whether these genomic features are reflected at the population level, we complemented the reference-based analysis with a reference-independent k-mer approach using WGS from 174 wild oysters (88 large, 86 small). We identified three distinct classes of sequence variation differentiating the cohorts: (1) bacterial DNA sequences enriched in smaller oysters; (2) a host genomic signal localizing to the *RNF213* locus on Chromosome 1; and (3) low-complexity repetitive elements enriched in larger oysters. These findings support the expansions seen in the assembly of the reference genome and lends evidence for the dynamic nature of the oyster genome. The data points to the 13Mb region as not just representative of environmental-related evolutionary divergence among species, but indicates that this region is a sort of genomic “neighborhood” [48, 49] that actively varies in wild populations of oysters.

The genome-wide association study (GWAS) of oyster body size provided additional population-level evidence linking this genomic neighborhood to phenotypic variation. We identified a SNP at position 35,698,255 on Chromosome 1 (A>G) that was significantly associated with the Large phenotype (Fisher’s exact test, p = 0.0002514). The G allele was present in 7 of 8 Large oysters but absent from all 10 Small oysters tested. Although this SNP resides within an intron of the uncharacterized gene *g2168*, its location within the 13 Mb immune-enriched region. Its strong association with size suggests that structural or regulatory variation in this neighborhood may influence growth and physiological performance in wild populations [50, 51]. The convergence of three independent analytical approaches on the same Chromosome 1 interval strengthens the conclusion that this region represents a functionally relevant genomic neighborhood shaped by environmental pressures [48–50].

The gene expansions noted in the reference genome are consistent with prior genomic studies in oysters that highlight both lineage-specific gene expansions and structural plasticity [27, 28]. Peng et al. (2020) generated a chromosome-level genome assembly for *C. hongkongensis* using a combination of long and short-read sequencing technologies and linkage mapping, achieving high completeness (∼95.7% BUSCO) [27]. Comparative analyses revealed strong conservation at the chromosomal level but substantial diversification in gene content, particularly through expansion of innate immune gene families such as Toll-like receptors (TLRs) [27]. Similarly, Powell et al. (2018), in their assembly of the *Saccostrea glomerata* genome, identified extensive expansion of transposable elements, including nucleases and transposases, suggesting that mobile genetic elements contribute to genome restructuring [28]. Our study’s data and conclusions agree with those of Peng et al. (2020) and Powell et al. (2018), and demonstrates that oyster genomes are both structurally dynamic and evolutionarily responsive. Both gene expansion and repeat-mediated remodeling appear to be acting as complementary mechanisms that facilitate species-specific adaptation to environmental exposures.

The expansion of immune-related gene families observed across oyster species likely reflects the unique ecological and physiological demands placed on these organisms. As sessile filter feeders, oysters are continuously exposed to a wide array of pathogens and environmental stressors, including fluctuations in temperature, salinity, and pollutant load [3–11]. In the absence of an adaptive immune system, this persistent exposure necessitates a robust innate immune response. Gene duplications and expansions may provide a mechanism for increasing both the diversity and responsiveness of immune signaling pathways, allowing oysters to detect and respond to diverse environmental challenges [27, 28]

The population-level k-mer analysis conducted in parallel with this genome assembly provides a direct illustration of these ecological pressures in action. By comparing oysters stratified by body size, we identified a striking enrichment of bacterial DNA sequences from the *Spirochaetota* and *Actinomycetota* phyla [80] in the smaller cohort. These bacterial sequences were detectable only through reference-independent k-mer analysis and were absent from all assembled oyster reference genomes [2, 75], highlighting the capacity of sentinel organisms to concentrate and retain molecular evidence of environmental exposures [20]. While the presence of microbial DNA does not establish causation, smaller oysters may be more susceptible to infection, or infection may impair growth [81]. This finding underscores the reality that wild oyster populations continuously interface with a diverse microbial exposome [3–11].

The 13 Mb immune neighborhood on Chromosome 1, characterized by expansions in cytokine-signaling and stress-response genes, likely represents the host genomic architecture that has evolved to meet this persistent pathogenic pressure.

### Contextualizing the Evolutionary Duplications Observed in the Reference Genome

Oysters’ constant exposure to the environment creates their uniqueness in understanding genomic rearrangements. One example of such a fluctuation that could have prompted evolutionary adaptation in *C. virginica* could be rising ocean temperatures. Since 1901, US coastal Atlantic ocean surface temperatures have risen, on average, approximately 0.072°C per year, with a rapid increase (approximately 1.5°C overall) since 1980 [29, 30]. This means that the environmental temperature for *C. virginica* has continually been getting warmer over the past century, causing a need to cope with this new stress. These environmental changes may contribute to the selective pressures that favor the *C. virginica* genome restructuring for gene expansions on chromosome 1 to upregulate the activity of immune and stress related functions. In particular, we found many of these genes to be linked to cytokine-signaling pathways. While the expansions we observed likely reflect selective pressures accumulated over evolutionary timescales rather than solely the past century, the sustained directional trend in coastal temperatures illustrates the type of chronic environmental stress that could favor expanded immune capacity over generational timescales.

In the context of rising ocean temperatures, one possible mechanistic explanation for the expansion of cytokine-associated genes involves heat shock proteins (HSPs). HSPs proteins are highly conserved across phylogeny, spanning both prokaryotic and eukaryotic organisms [31], and play a fundamental role in maintaining cellular homeostasis under stress conditions [32]. Two decades worth of literature as contributed to understanding HSPs as innate immune system activators, such as those of cytokine-induced proinflammatory responses [33–40]. HSPs have been implicated in immune signaling processes, including modulation of cytokine production, enhancement of antigen presentation, and functioning as molecular markers that alert the immune system to cellular damage [32]. Although some studies have raised concerns about experimental procedures in earlier work—particularly regarding the purity of HSP preparations and their direct role in immune activation [41]—the broader conceptual link between cellular stress responses and immune signaling remains well supported.

However, HSPs are just one instance in the face of one environmental exposure. More importantly, one should note the coordinated regulation of genes that experience duplication events within non-conserved contiguous regions. We see this clearly with Peng et al. (2020), with their reference genome indicating high levels of duplications for genes involved with TLRs.

In the innate immune system, dendritic cells, granulocytes, and macrophages protect against pathogens using TLRs that trigger intracellular signaling cascades, activating transcription factors such as NF-κB [42]. This promotes the expression of pro-inflammatory cytokines and other immune mediators [42]. TLRs have been found to be conserved across phylogeny for both vertebrates and invertebrates [43, 44]. In organisms such as oysters, which lack adaptive immunity, TLR-mediated signaling plays an especially critical role in maintaining immune competence [45, 46]. As sessile organisms continually exposed to microorganisms in their environment, TLRs serve as a prominent backbone of the mollusk immune system [45, 46]. This provides context for the duplications of genes observed in the study by Peng et al and lends further perspective to the expansion of cytokine-related genes in *C. virginica*.

Importantly, TLR signaling is not a simple linear pathway but a highly regulated network in which signal strength, duration, and downstream transcriptional responses are finely tuned through multiple layers of control, including chromatin remodeling, transcription factor recruitment, and post-transcriptional regulation of mRNA stability [47]. To directly assess the transcriptional responsiveness of the expanded region we identified, we analyzed RNA-seq data from oysters challenged with 2′3′-cUA, a cyclic dinucleotide that activates the innate immune system [23–25]. Within the 13 Mb region, 87 genes showed at least 2-fold upregulation following 2′3′-cUA treatment. This coordinated regulatory activation indicates that this structurally dynamic genomic region is not merely expanded but also transcriptionally responsive to immune stimulation. These findings provide functional support for the hypothesis that expansion of this region enhances the capacity for rapid and coordinated immune responses.

In this context, the spatial clustering of immune- and stress-related genes within expansion-rich regions—such as the ∼13 Mb region identified on chromosome 1 of *C. virginica*—may facilitate coordinated gene regulation, enabling more efficient and synchronized transcriptional responses to environmental stimuli. Such genomic neighborhoods could enhance the speed and precision with which immune pathways are activated, particularly under conditions of sustained or repeated stress exposure.

This model of thinking is only one possible explanation for the observed duplications seen in *C. virginica* and other oyster species. It should not be taken as an absolute theory, as these mechanisms are inferred from conserved biological systems rather than directly demonstrated in oysters beyond genomic architecture. However, they do provide a valuable comparative framework for understanding gene-environment interactions more broadly. We highlight the fact that ambient environmental stressors, like rising temperatures, play a key role in modulating the regulation of immune and stress response genes. Stress-response proteins seem to act as key interfaces between environmental exposure and immune activation. In oysters, which experience constant and often extreme environmental variability, expansions of such gene families may therefore confer an adaptive advantage by increasing the sensitivity, specificity, and scalability of stress detection and response systems. The shared evolutionary ancestry of stress-response pathways across species suggests that insights gained from oyster genomic adaptation may inform our understanding of how environmental pressures influence immune and stress-related processes in humans.

### Reference-Independent Evidence Reveals Environmental and Microbial Dimensions of Genomic Variation

Above presents molecular and evolutionary arguments that gene duplications within immune-enriched genomic regions confer adaptive advantages under chronic environmental stress. These findings are further supported by the population-level evidence derived from reference-independent k-mer analysis conducted in parallel with this genome assembly. As described earlier, oysters stratified by body size exhibited significant differences in both microbial sequence signatures and host genomic variation within the 13 Mb immune-enriched region on chromosome 1.

Specifically, smaller oysters exhibited enrichment of non-reference bacterial DNA sequences, including signatures from the *Spirochaetota* and *Actinomycetota* phyla [80]. These findings are consistent with increased microbial exposure, altered host-microbe interactions, or differences in immune clearance, although the directionality of this relationship cannot be determined from the present data [81]. However, sequencing data from the small cohort of oysters demonstrated enrichment of k-mers mapping to the *RNF213* locus, a gene located within the chromosome 1 immune neighborhood and encoding an E3 ubiquitin-protein ligase implicated in stress-response and immune-related pathways [82, 83]. Together, the co-occurrence of microbial sequence enrichment and host genomic variation, and the localization of this host signal to a region enriched for lineage-specific immune gene expansion, point to population-level evidence that this genomic neighborhood is functionally relevant in shaping responses to environmental stress.

While the HSP and TLR mechanisms discussed above offer plausible molecular explanations for the evolutionary expansion of immune-related genes, the k-mer findings extend this framework by demonstrating that these genomic features are associated with variation observed in contemporary wild populations [48–50]. Rather than representing solely historical adaptations, the 13 Mb region on chromosome 1 appears to function as a dynamic genomic landscape in which environmental exposures, microbial interactions, and host genomic variation intersect [48, 49]. Importantly, these findings also highlight the value of integrating reference-independent approaches with traditional genome assembly and annotation. The bacterial sequences identified in this analysis were not captured in reference genomes and would likely have been missed using alignment-based methods alone [84]. Similarly, the identification of host-derived variation within the *RNF213* locus through k-mer analysis accentuates the ability of sequence-based approaches to detect signals that may not be readily apparent through SNP-focused analyses [58–60].

The GWAS-identified SNP at HiC_scaffold_1:35,698,255 localizes to the same 13 Mb Chromosome 1 region that harbors RNF213 and the lineage-specific immune gene expansions identified in our reference assembly. While the k-mer analysis captured sequence-level variation at RNF213 enriched in Small oysters, the GWAS identified a common SNP associated with the Large phenotype. Rather than representing conflicting signals, these findings suggest that multiple classes of genomic variation—including single nucleotide polymorphisms, structural variation, and repetitive elements—coexist within this dynamic region and may differentially associate with physiological outcomes [48–50, 52, 55].

One possible interpretation is that variation within this genomic neighborhood reflects differences in environmental exposures experienced by wild oyster populations [3–11, 67–69].

Smaller oysters, which exhibited enrichment of microbial sequences, may be consistent with increased exposure to pathogens or environmental stressors, potentially interacting with underlying genomic variation to influence growth and physiological performance [3–11, 19]. In this context, the Chromosome 1 region may function as a dynamic genomic interface through which environmental pressures and host genetic variation jointly shape phenotype, rather than acting through single variants in isolation [48–50, 52, 55, 67–69].

Taken together, the convergence of evolutionary genomics, functional annotation, and population-level k-mer sequence analysis on the same genomic region strengthens the argument that immune-enriched genomic neighborhoods represent a fundamental unit of adaptation to environmental stress [48–50, 52, 55]. In this framework, adaptation is not driven solely by individual variants, but by coordinated changes across clusters of functionally related genes embedded within dynamic genomic regions [52, 55, 57, 61]. This perspective has direct implications for chronic disease epidemiology, where complex, long-term environmental exposures may similarly influence disease risk through coordinated genomic responses rather than isolated genetic effects [67–69].

### C. Virginica as a Sentinel Species for Understanding Population-Level Changes in Genomic Neighborhoods in Response to Environmental Exposures

The findings of this study present a high quality reference genome and genomic analysis that can be used to better understand how genome architecture responds to sustained environmental stress. Unlike many terrestrial organisms, oysters cannot behaviorally avoid adverse conditions, meaning that adaptation must occur at the physiological and genomic levels [48]. This makes them particularly well-suited for studying how persistent environmental pressures shape gene expansion, regulatory architecture, and immune responsiveness over evolutionary timescales. While oysters lack an adaptive immune system, rendering them useful only as sentinels rather than direct model organisms, the genome *C. virginica* and its comparative analyses to other species offers the chance to understand the restructuring of the genome in response to environmental exposures.

The host-genomic signal detected in our k-mer analysis provides support for this sentinel framework. Among the k-mers significantly enriched in the Small cohort, a subset mapped unambiguously to the *C. virginica* host genome, localizing specifically to *RNF213* on Chromosome 1. *RNF213* encodes an E3 ubiquitin-protein ligase implicated in innate immune signaling and cellular stress responses [82, 83], and it resides within the 13 Mb immune-enriched genomic neighborhood characterized in our reference assembly [75]. The association between variation at this locus and physiological condition in a wild population demonstrates that this genomic neighborhood is not merely a static feature of the oyster genome but is actively varying in response to real-world environmental pressures. This finding supports the sentinel species concept: the oyster genome contains identifiable, measurable features that reflect the cumulative burden of environmental exposures at the population level.

The GWAS findings extend this sentinel framework in an important direction. While the k-mer analysis identified sequence variation at RNF213 associated with the Small phenotype, the GWAS identified a common SNP in a nearby gene (*g2168*) strongly associated with the Large phenotype. This bidirectional pattern—distinct classes of genomic variation associating with opposite phenotypic extremes within the same genomic neighborhood—suggests that this region may harbor a suite of variants influencing growth and immune competence [48–50, 52, 55]. In human populations, analogous genomic neighborhoods may similarly contain multiple variants that collectively modulate disease susceptibility, only a subset of which are captured by standard SNP-based GWAS [58–60]. The oyster system thus demonstrates that fully characterizing complex traits requires integrating multiple forms of genomic variation within biologically meaningful regions, rather than considering variants in isolation [48–50, 58–60].

Furthermore, *RNF213* has direct implications for public health. In human populations, genome-wide association studies point to *RNF213* as the gene linked to Moyamoya disease (MMD), a chronic cerebrovascular disorder characterized by progressive stenosis of the internal arteries that often lead to fatal stroke [85, 86]. MMD is approximately 10-fold more prevalent in East Asian populations than in Western countries, and the mutation (p.R4859K in *RNF213*) is present in approximately 95% of Japanese familial cases, 73% of non-familial cases, and 1.4% of unaffected controls (OR = 190.8) [85]. Notably, this variant is largely absent in Caucasian controls and Caucasian MMD patients [86]. While MMD exhibits familial clustering, the data clearly indicate that genetics alone cannot explain disease onset. The p.R4859K variant demonstrates incomplete penetrance, with many carriers remaining asymptomatic throughout life, and homozygous carriers have been identified among healthy controls [85, 86]. This pattern strongly implicates gene-environment interactions in disease pathogenesis. Notably, *RNF213* is highly expressed in immune tissues, particularly the spleen and leukocytes, rather than in the brain itself [85, 86]. Functional studies have demonstrated that *RNF213* is upregulated by inflammatory signals, including interferons and cytokines, and plays a role in cell-autonomous immunity against intracellular pathogens [86].

MMD illustrates a broader principle:susceptibility to complex disease often resides not in single nucleotide variants acting in isolation, but in the coordinated behavior of genomic neighborhoods. The p.R4859K variant in *RNF213* confers extraordinary risk (OR = 190.8) [85], yet the majority of carriers never develop disease. This incomplete penetrance indicates that the variant alone is insufficient. Disease manifests only when the genomic region in which *RNF213* is embedded encounters specific environmental triggers, such as infection, inflammation, or hemodynamic stress. However, this cause has not been fully resolved [85, 86].

Critically, if researchers had approached MMD using only the standard tools of human genomics, such as searching for SNPs associated with disease status while controlling for environmental covariates, the essential gene-environment interaction would have been obscured.

This perspective has direct implications for chronic disease epidemiology. Complex diseases such as cardiovascular disease, autoimmune disorders, and metabolic syndrome are increasingly understood to arise from the interplay of polygenic susceptibility and cumulative environmental exposures [67, 68, 87]. Yet the dominant analytical paradigm in human genomics, the GWAS, treats the genome as a collection of independent SNPs and the environment as a set of covariates to be controlled [59, 60]. The sentinel framework advanced here suggests an alternative approach: identifying and monitoring genomic neighborhoods that are structurally variable across populations and transcriptionally responsive to environmental challenge.

In the oyster, the 13 Mb region on Chromosome 1 is such a neighborhood—expanded in a lineage-specific manner, enriched for immune genes, upregulated following immune stimulation, and varying with physiological condition in wild populations. In humans, syntenic regions may function as analogous neighborhoods. Expanding our understanding of genetic disease beyond the sequencing and analysis of isolated variants to include the structural and regulatory architecture of these neighborhoods will be essential for capturing the full complexity of gene-environment interactions [62, 63, 65, 66]. The oyster, by concentrating environmental exposures and making their genomic consequences measurable, provides both the proof of principle and the methodological roadmap for this expanded vision [52, 53, 55, 57].

Just as different species of oysters have different ambient stressors, different populations of humans will experience an uneven distribution of stressors across sub populations. The presence of genomic neighborhoods may allow species to tailor their immune and stress responses through coordinated genomic regulation [51]. As such, it follows that similar genomic neighborhoods may be observed in humans. In fact, the literature supports such a model. In humans, genes with similar expression patterns are often co-expressed within shared chromatin regions, allowing for the regulation of genes not just at the individual level, but regionally [52–55]. This provides the physical genomic architecture that allows for genomic neighborhoods to develop. The gene neighborhoods often arise from tandem duplication events, suggesting that they are not just co-expressed, but functionally related [54, 56, 57]. Thus, it seems that both humans and oysters share the capacity for their genomes to have tandem duplication of genes, representing a shared evolutionary trait of their genomes.

Translating this framework to human studies would shift how we investigate the health impacts of chronic environmental exposures. Firstly, using a genomic neighborhood framework offers to better identify structural variants and copy number variations than current analyses. The literature has pointed out that structural variants effect more base pairs than SNPs [62], and it is well known that SNP arrays are limited in their power to detect copy number variations [63]. In the context of innate immunity, it seems that genes involved in the regulation of stress and immune responses are clustered as coordinated units rather than collections of independent variants [64], which is also clearly demonstrated in the genome of *C. virginica*. Furthermore, it has been suggested that environmental exposures can alter 3D chromatin structure, affecting multiple genes simultaneously [65, 66]. Thus, by only analyzing SNPs, we are losing the complexity of what is actually happening at the chromosomal level, and there is a need to shift frameworks from SNPs to genomic neighborhoods to better understand the association between the genome and environmental stressors. The genomic neighborhoods framework is not a replacement for SNP-based studies, which remain valuable for their scalability and well-established statistical methods. Rather, it represents a complementary approach for research questions that SNPs cannot fully address.

This could advance our understanding of the genome in the face of the social determinants of health and their correlation with the onset of chronic conditions. Environmental exposures, like climate change or socioeconomic status, are complex, multifactorial, and unevenly distributed across populations [67]. These exposures are unlikely to influence health outcomes through isolated genetic variants alone, but rather through coordinated changes in gene regulation, chromatin architecture, and genomic organization [68]. A genomic neighborhood framework provides a more biologically realistic model for capturing these interactions, as it accounts for how groups of functionally related genes may respond collectively to sustained environmental pressures [69]. In this way, oysters serve as a sentinel system that allows us to observe how chronic exposure shapes genome architecture in real time, offering insight into the mechanisms by which environmental stressors may contribute to population-level differences in immune function and disease susceptibility in humans [48].

### Limitations and Future Direction of Research

Several limitations of this study warrant consideration and suggest avenues for future research. First, the cross-sectional design of the K-mer association analysis precludes causal inference. We cannot determine whether bacterial burden preceded and contributed to reduced growth, or whether smaller oysters were more susceptible to infection due to pre-existing physiological compromise. Longitudinal sampling of individual oysters across seasons, coupled with repeated measurements of growth, immune markers, and microbial burden, would help resolve directionality.

Furthermore, this study only sampled oysters from a small geographic region on the coast of the Easter United States. While this has provided an excellent foundation from which to resolve the genome assembly of *C. virginica*, expanding sampling to populations across the Atlantic coast and Gulf of Mexico, which experience different thermal regimes, salinity profiles, and pathogen communities, would reveal whether the 13 Mb immune neighborhood varies along environmental gradients in predictable ways.

The independent validation cohort was limited to 17 samples (8 Large, 9 Small). While the direction of effect was consistent with the discovery analysis, the difference did not reach statistical significance (p = 0.577). With this sample size, the analysis had limited statistical power to detect modest effect sizes, and a substantially larger validation cohort would be required to confirm the population-level signal with confidence. Additionally, we suggest using varying lengths of k-mers in future analyses to capture larger changes in the genome that may have been occluded by our set lengths.

Finally, when extrapolating information learned from oysters to humans, one must account for the difference in their immune systems. While oysters can’t be used as model organisms for human health, this paper does not claim to use oysters as such. Rather, oysters are sentinels, and serve to show lessons learned in the gene-environment interaction and how environmental exposures influence our genome and lead to chronic conditions. Here, we have developed a framework from which to think about how to better understand genomic changes as “neighborhoods” that lead to coordinated regulatory features in response to exposures. However, while the human *RNF213*-Moyamoya connection provides a compelling cross-species parallel, direct functional validation of *RNF213* in oysters is lacking. Future studies employing gene knockdown (RNAi) or CRISPR-based approaches in oyster hemocytes or larvae would be valuable for establishing the mechanistic role of *RNF213* in oysters.

The GWAS component of this study also has important limitations. The sample size for initial SNP discovery was modest (n = 18), and the identified SNP (p = 0.0002514) does not meet genome-wide significance thresholds after multiple-testing correction. Validation in larger, independent cohorts is essential to confirm this association and to determine whether the variant itself, or linked functional variants, underlie the observed size differences. Additionally, because the SNP lies within a non-coding intron of an uncharacterized gene, functional studies—including expression quantitative trait locus (eQTL) analysis, chromatin accessibility assays, or CRISPR-based perturbation—will be required to establish mechanistic links between this locus and growth phenotypes. Finally, expanding sampling to populations across the Atlantic and Gulf coasts will reveal whether the frequency of this variant varies along environmental gradients in predictable ways.

## Conclusion

This study demonstrates that chronic environmental exposure leaves measurable signatures in the structural organization of the genome. By generating a high-quality reference assembly for *Crassostrea virginica*, conducting a genome-wide association study of body size, and complementing these approaches with reference-independent k-mer analysis, we identified a dynamic genomic neighborhood on chromosome 1 enriched for immune and stress-response genes that varies with physiological condition in wild populations. Three independent analytical methods converged on this same genomic interval, revealing multiple classes of variation including lineage-specific gene expansions, a size-associated SNP, and host sequence variation at the *RNF213* locus. These findings establish oysters as a sentinel system for observing how sustained environmental pressures shape genome architecture. More broadly, this work argues for expanding our analytical frameworks, from isolated genetic variants to coordinated genomic neighborhoods, to better capture the complexity of gene-environment interactions in both environmental sentinels and human populations.

## Acknowledgements

We thank the team at Oyster Farming Technologies for their support, patience, and knowledge of oysters and their ecology.

This manuscript is based on Walker Bargmann’s MPH thesis, for which he received the Yale School of Public Health’s 2026 Dean’s Prize for Outstanding MPH Thesis.

